# Tools and best practices for allelic expression analysis

**DOI:** 10.1101/016097

**Authors:** Stephane E. Castel, Ami Levy Moonshine, Pejman Mohammadi, Eric Banks, Tuuli Lappalainen

## Abstract

Allelic expression (AE) analysis has become an important tool for integrating genome and transcriptome data to characterize various biological phenomena such as *cis*-regulatory variation and nonsense-mediated decay. In this paper, we systematically analyze the properties of AE read count data and technical sources of error, such as low-quality or double-counted RNA-seq reads, genotyping errors, allelic mapping bias, and technical covariates due to sample preparation and sequencing, and variation in total read depth. We provide guidelines for correcting and filtering for such errors, and show that the resulting AE data has extremely low technical noise. Finally, we introduce novel software for high-throughput production of AE data from RNA-sequencing data, implemented in the GATK framework. These improved tools and best practices for AE analysis yield higher quality AE data by reducing technical bias. This provides a practical framework for wider adoption of AE analysis by the genomics community.

## INTRODUCTION

Integrating genome and transcriptome data has become a widespread approach for understanding genome function. Allelic expression (AE; also called allele-specific expression or allelic imbalance) analysis is becoming an increasingly important tool for this, as it quantifies expression variation between the two haplotypes of a diploid individual distinguished by heterozygous sites (Fig. 1a). This approach can be used to capture many biological phenomena (Fig. 1b): effects of genetic regulatory variants in *cis*^1–8^, nonsense-mediated decay triggered by variants causing a premature stop codon^9–12^, and imprinting^13, 14^. Standard RNA-sequencing data captures allelic expression only when higher expression of one parental allele is shared between individual cells (Fig. S1), as opposed to random monoallelic expression of single cells that typically cancels out when a pool of polyclonal cells is analyzed^15, 16^.

**Figure 1.**
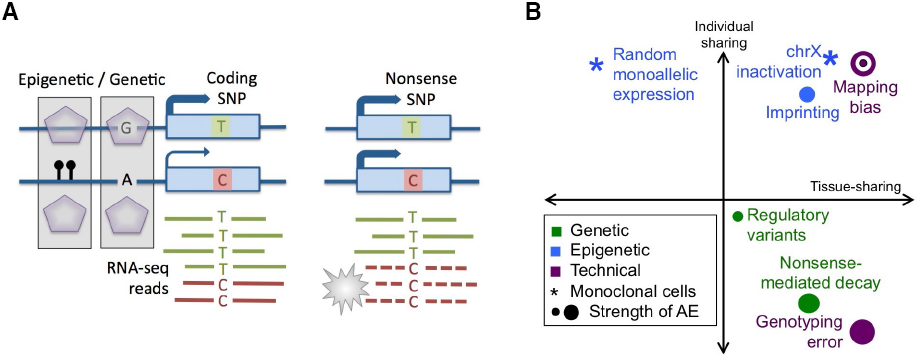
Allelic expression and its sources. A) Schematic illustration of allelic expression; B) Biological sources of AE, with the x-axis denoting the approximate sharing of AE across tissues of an individual, and the y-axis having the estimated sharing of AE signal in one tissue across different individuals^5, 8, 12, 13, 15^.

In this paper, we describe a new tool in the Genome Analyzer Toolkit (GATK) software package for efficient retrieval of raw allelic count data from RNA-sequencing data, and analyze the properties of AE data and the sources of errors and technical variation, with suggested guidelines for accounting for them. While most types of errors may be rare, they are easily enriched among sites with allelic imbalance, and can sometimes mimic the biological signal of interest, thus warranting careful analysis. Our focus is on methods for obtaining accurate data of allelic expression rather than building a GUI pipeline^17^ or downstream statistical analysis of its biological sources^9, 13, 18–20^. The example data in most of our analysis is the open-access RNA-sequencing data set of the LCLs of 1000 Genomes individuals from the Geuvadis project^5^. The scripts, tools and data are available in https://www.broadinstitute.org/gatk/ and https://tllab.org.

## RESULTS

### Unit of allelic expression data

The biological signal of interest in allelic expression analysis is the relative expression of a given transcript from the two parental chromosomes. Typical AE data seeks to capture this by counts of RNA-seq reads carrying reference and alternative alleles over heterozygous sites in an individual (het-SNPs), and this is the focus of our analysis unless mentioned otherwise. The Geuvadis samples with a median depth of 55 million mapped reads have about 5,000 het-SNPs covered by ≥30 RNA-seq reads, distributed across about 3,000 genes and 12,000 exons (Fig. 2, Fig. S2). The exact number varies due to differences in sequencing depth, its distribution across genes, and individual DNA heterozygozity. About one half of these genes contain multiple het-SNPs per individual, which could be aggregated to better detect allelic expression across the gene (Fig. 2d). However, alternative splicing can introduce true biological variation in AE in different exons, and incorrect phasing needs to be accounted for in downstream analysis^13^. Additionally, summing up data from multiple SNPs is not appropriate if the same RNA-sequencing reads overlap both sites. In the Geuvadis data, 9% of the reads used in AE analysis in fact overlap more than one het-SNP (Fig S2d), but this will become more frequent as read lengths increase^21^. In the future, better tools are needed to partition RNA-seq reads to either of the two haplotypes according to all het-SNPs that they overlap^22^. In fact, this could help to phase exonic sites separated by long introns.

**Figure 2.**
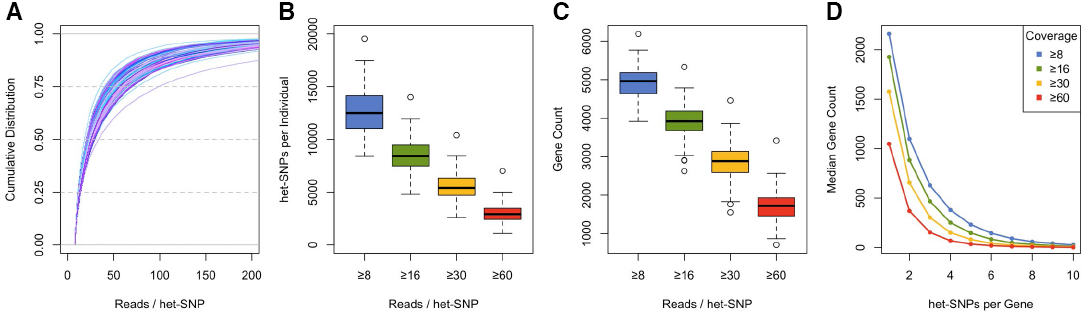
Genomic coverage of allelic expression data in Geuvadis CEU samples. A) Cumulative distribution of RNA-seq read coverage per het-SNP (each line represents one sample). B-C) The number of het-SNPs (b) and protein-coding genes (c) per sample as a function of coverage cutoff. D) The number of protein-coding genes with AE data vs the number of het-SNPs they contain. Each line is the median for all samples at a specific coverage level.

AE analysis of small insertions or deletions (indels) has proven to be technically very challenging and it is rarely attempted even though frameshift indels are an important class of protein-truncating variants. Alignment errors over indel loci are pervasive due to multiple mismatches of reads carrying alternative alleles, and lower genotyping quality adds further error^12^. In Rivas *et al*. we describe the first approach for large-scale analysis of allelic expression over indels, but further methods development is warranted for better sensitivity and computational scalability.

In addition to classical AE analysis to detect differences in total expression level of two haplotypes, it is also possible to analyze allelic differences in transcript structure or splicing (AS;^5, 21^). These methods compare the exon distribution of reads and their mates carrying different alleles of a heterozygous site, and work increasingly well for longer total fragment lengths. In these analyses, the data structure is somewhat more complex than reference / non-reference read counts in AE, depending on the specific algorithm. While this paper focuses on classical AE analysis of SNPs, most of the quality analysis steps apply to indel AE and AS analyses as well.

### Tools to retrieve allele counts

To retrieve allele counts we have developed a new GATK tool, named ASEReadCounter, which will be publicly available in the GATK version 3.4. The GATK^23, 24^, is a software package that offers a wide variety of tools to analyze high-throughput sequencing data. It is also a programming framework that allows easy development of new genome analysis tools such as the one detailed in this section.

The ASEReadCounter tool operates on aligned RNA-seq reads and counts the alleles over heterozygous sites. For each bi-allelic site (multi-allelic sites are currently ignored) the tool counts the reference and alternative allele bases that passed filters for mapping and base quality. ASEReadCounter offers several options for processing RNA-sequencing reads: by default each read fragment is counted only once if the base calls are consistent at the site of interest, and duplicate reads are filtered (see below). Additional options allow filtering for coverage and for sites or reads with deletions. The output of the new tool is one file per RNA-seq input file, with one line per site displaying the counts for each allele as well as counts of filtered reads. Several output formats are available, and the default output file is compatible with downstream tools, such as the statistical analysis tool MAMBA^20^.

### Quality control of allele counting

Retrieving allele counts from RNA-seq data over a list of heterozygous sites is conceptually very simple, but several non-trivial filtering steps need to be undertaken to ensure that only high-quality reads representing independent RNA/cDNA molecules are counted. The first commonly applied filter is to remove reads with a potentially erroneous base over the heterozygous site based on low base quality. Furthermore, potential overlap of mates in paired-end RNA-sequencing data needs to be accounted for, so that each fragment, representing one RNA molecule, is counted only once per het-SNP. In the Geuvadis data, an average of 4.4% of reads mapping to het-SNPs per sample are derived from overlapping mates, but this number will vary by the insert size (Fig. S3a).

In RNA-seq analysis, duplicate reads with identical start and end position are common (15% of reads in Geuvadis AE analysis), because highly expressed genes get saturated with reads (Fig. S3b–d). Thus, by default duplicates are usually not removed from RNA-seq data to avoid underestimating expression levels in highly expressed genes^5^. However, we observe consistent albeit infrequent signs of PCR artifacts in the Geuvadis AE data, affecting especially lowly covered sites – where duplicates are mostly true PCR duplicates, since saturation is unlikely. Removing duplicate reads reduces technical sources of AE at these sites, while having a minimal effect on highly covered, read saturated SNPs (Fig. S3e). Thus, we suggest that removing duplicate reads is a good default approach for AE analysis, and it is implemented as a default in the GATK tool.

The most difficult problem in AE analysis and a potential source of false positive AE is ensuring that 1) all the reads counted over a site indeed originate from that genomic locus, and 2) all reads from that locus are counted. RNA-seq studies with shorter or single-end RNA-seq reads are more susceptible to these problems. First, to make sure that no alien reads get erroneously assigned to a locus, only uniquely mapping reads should be used. This implies that highly homologous loci - such as miRNAs - are not amenable to AE analysis.

An even more difficult caveat in AE analysis is allelic mapping bias: in RNA-seq data aligned to the reference genome, a read carrying the alternative allele of a variant has at least one mismatch, and thus has a lower probability to align correctly than the reference reads^25^-^27^. Simulated data in Panousis et al.^26^ indicates substantial variation between sites - in most cases reads mapped correctly, but 12% of SNPs and 46% of indels had allele ratio bias >5% with some having a full loss of mapping of the alternative allele. Loci with homology elsewhere in the genome are particularly problematic as reads have nearly equally good alternative loci to align to. Furthermore, even a site with no bias in itself can become biased due to a flanking (sometimes unknown) variant that shares overlapping reads with the site of interest. In addition, mapping bias varies depending on the specific alignment software used (Fig. S4a–c).

Various strategies can be employed to control for the effect of mapping bias on AE analysis. The simplest approach that can be applied to AE data without realignment is to filter sites with likely bias^5,8,27^. In previous work^5,8,28–30^ and in this paper unless mentioned otherwise, we remove about 20% of het-SNPs that either fall within regions of low mappability (ENCODE 50bp mappability score < 1) or show mapping bias in simulations^26^. This reduces the number of sites with strong bias by about 50% (Fig. 3a–c), but the genome-wide reference ratio remaining slightly above 0.5 indicates residual bias (Fig. S5a). Using this ratio as a null in statistical tests instead of 0.5^5,6^ can improve results (Fig. S5b–e). A more exhaustive but computationally intensive approach is alignment to personalized genomes^18,31,32^ or use of a variant-aware aligner, such as mrsFAST-Ultra^33^, and GSNAP^34^. These methods yield comparable results and eliminate *average* genome-wide bias (Fig. 3d–f, S4d–f), but mappability filters are still essential to remove at least most of the sites where homology elsewhere in the genome leads to substantial allelic mapping bias (Fig. S4g–i). Altogether, while many commonly used approaches yield reasonably accurate data, allelic mapping bias remains a problem without a perfect solution.

**Figure 3.**
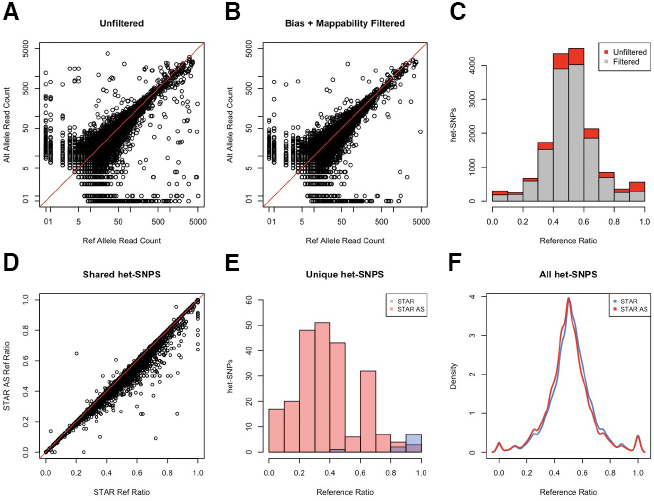
Strategies for reducing mapping bias in allelic expression analysis. A-C) AE data using a standard alignment approach (STAR 2-pass) before (a) and after (b) filtering sites based on low mappability and simulated mapping bias, and the resulting reference allele ratios (c). D-F) The effect of mapping to a personalized reference on AE data, with reads aligned using STAR to either hg19 (STAR, blue) or a personalized genome generated using AlleleSeq (STAR AS, red). Scatterplot of reference ratio at sites with AE data in both mapping strategies (shared het-SNPs, d), histogram of reference ratios at sites with AE data in only one mapping strategy (unique het-SNPs, e), and overall distribution of reference ratios using each mapping strategy (all het-SNPs, f). Sites with low mappability and simulated mapping bias have been excluded from d-e.

### Quality control of genotype data

AE analysis relies on data of heterozygous sites to distinguish the two parental alleles. These genotype data are ideally retrieved from DNA-sequencing or genotyping arrays, but the RNA-seq data itself can also be used for calling genetic variants and finding heterozygous sites (^35,36^, https://www.broadinstitute.org/gatk/guide/best-practices). However, true allelic imbalance can lead to heterozygous sites being called homozygous in RNA-based genotype calling and lead to substantial error in monoallelic genes due to e.g. imprinting, and more subtle bias in eQTL genes (Fig. S6a).

Even when using heterozygous genotypes called from DNA data, genotyping error can be an important source of false signals of allelic imbalance, because AE data from a homozygous site appears as monoallelically expressed. In genotype data that has passed normal quality control including Hardy-Weinberg equilibrium test, genotype error will lead to rare cases of monoallelic expression per site, not shared across many individuals (Fig. 1b). False heterozygous genotype calls are rare but not negligible in AE analysis using SNP genotypes from arrays or from modern sequencing data, but much more common in imputed data. Calculating the genome-wide proportion of monoallelic AE sites per individual is a sensitive method for genotyping quality control.

Removing genotyping error is relatively straightforward for analysis of moderate allelic imbalance (such as that caused by *cis*-regulatory variants): removing monoallelic variants removes sites with false genotypes and results in little loss of truly interesting data. However, highly covered sites are rarely strictly monoallelic even in a homozygous state due to rare errors in sequencing and alignment (Fig. S6b). Thus, we propose a genotype error filter where the average amount of such sequencing noise per sample is first estimated from alleles other than reference (REF) or alternative (ALT) (Fig. S6c). Then, binomial testing is used to estimate if the counts of REF/ALT alleles are significantly higher than this noise, and sites where homozygosity cannot be thus rejected are flagged as possible errors (Fig. 4). Additionally, it may be desirable to flag fully monoallelic sites with low total counts, where homozygosity cannot be significantly rejected, but heterozygosity is not supported either. This test can also be applied to study designs with RNA-seq data from multiple samples (e.g. tissues or treatments) of a given individual, genotyped only once, since genotyping error causes consistent monoallelic expression in every tissue. In the Geuvadis data set with 1000 Genomes Phase 1 genotypes and sites covered by ≥ 8 reads, an average of 4.3% of sites per sample are excluded by these criteria (1% FDR).

**Figure 4.**
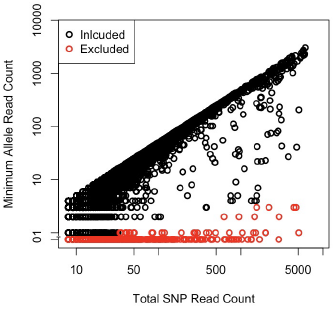
Quality control of genotype data for allelic expression analysis. Total het-SNP read count vs the read count of the lesser-covered allele for an individual Geuvadis sample. Sites flagged as putative genotyping errors are marked in red, with RNA-seq data not rendering support for heterozygosity.

Unfortunately, genotyping error is very difficult to distinguish from a true biological pattern of strong monoallelic expression, shared across all studied tissues, and present in a small number of samples, such as analysis of nonsense-mediated decay triggered by a rare variant, or a rare severe regulatory mutation (Fig. 1). The only real solution is rigorous genotype quality control and/or validation, and taking the possibility of confounding by genotyping error into account in interpretation of the results.

Sample mislabeling or mixing of the RNA-seq samples can lead to a substantial number false positive hits – as opposed to reduction of power in eQTL studies. Fortunately, simple metrics from AE analysis provide a sensitive way to detect sample contamination and mislabeling^37^. DNA-RNA heterozygous concordance – i.e. the proportion of DNA-heterozygous sites that are heterozygous also in RNA data – and a measure of allelic imbalance detect outliers and indicate the type of error (Fig. S6d).

### Technical covariates

RNA-seq has become a mature and highly reproducible technique, but it is not immune to technical covariates such as the laboratory which experiments were performed in, aspects of library construction and complexity, and sequencing metrics^37^. Gene expression studies are particularly susceptible to these technical factors, because read counts *between* samples are compared. AE analysis has the advantage that only read counts *within* samples are compared (allele vs allele), which makes it less susceptible to technical artifacts. We analyzed the correlation of the proportion of significant AE sites (binomial test, nominal p<0.05) with various technical covariates in the Geuvadis data (Fig. 5a). In raw AE count data, we observe a high correlation with the library depth (unique reads; R^2^ = 0.24) – expectedly, since total read count of AE sites determines the statistical power to see significant effects (see below). In AE data corrected for variation in read counts by scaling the counts to 30, all technical correlations are very small and mostly non-significant, in stark contrast to gene expression level data that displays strong batch effects (Fig. 5b). Thus, when appropriate measures are taken, AE analysis is an extremely robust approach that suffers less from technical factors than gene expression studies.

**Figure 5.**
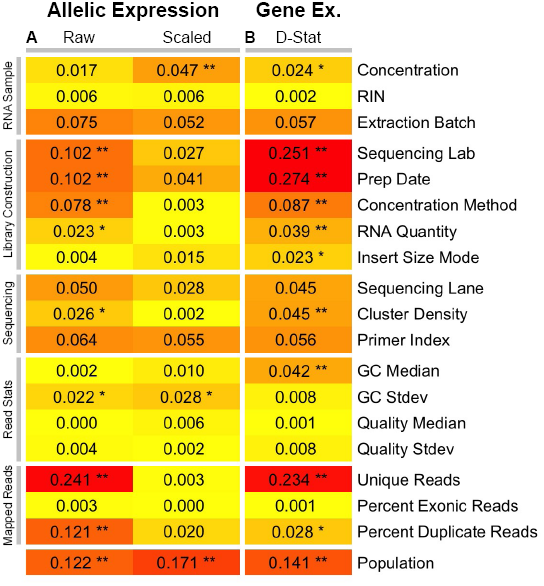
Technical covariates of allelic expression. A) Correlation of AE with technical covariates, measured as correlation (R^2^) between each covariate and the percentage of significant AE sites in a sample (binomial p < 0.05, het-SNPs with ≥ 30 reads), both before and after scaling to 30 reads. B) Correlation of gene expression with technical covariates. As the gene expression statistic we use the median correlation of each sample to all other samples (D-statistic). Correlation to a biological covariate (population) is shown for comparison. Correlations were calculated from all Geuvadis samples by Spearman Correlation for continuous covariates, or linear regression for categorical covariates. ** (p < 0.01), * (p < 0.05) after Bonferroni correction.

### Statistical Tests for Allelic Expression

A binomial test is the most classical way to test whether the ratio of the two alleles is significantly different from the expected 0.5. We show that after the QC measures outlined above, it provides a surprisingly good fit for the Geuvadis data (Fig. 6a–b, S7a), despite earlier reports suggesting substantial overdispersion^38, 39^. It does however consistently underestimate variance at both very lowly (<10) and very highly (>1000) covered sites, although QC measures reduce this. Overall, this renders strong support for the overall high quality of the allelic count data from modern RNA-seq experiments.

**Figure 6.**
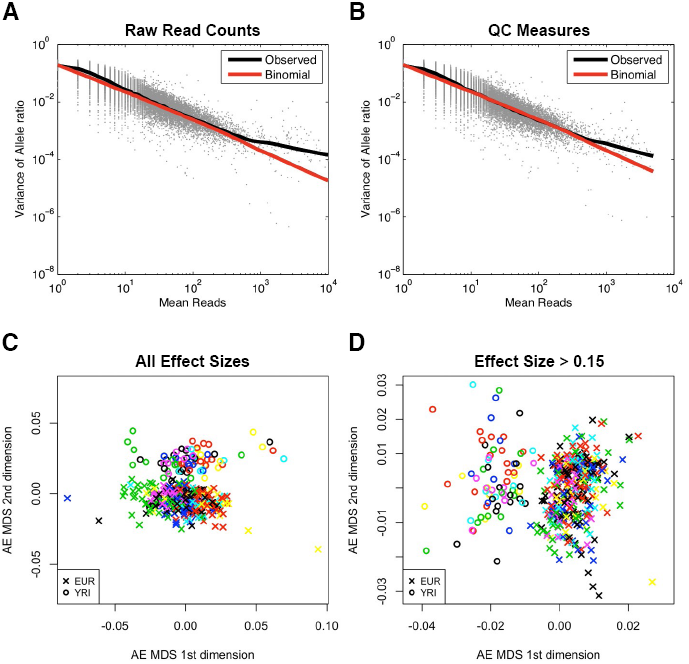
Binomial testing for significant allelic expression. A-B) Variance of allelic ratios as a function of total read counts, calculated as the mean at a given SNP from a Geuvadis individual with 8 technical replicates (grey) with (b) or without (a) QC filters. The lines denote locally weighted smoothing of observed data (black) and theoretical variance for binomially distributed data (red). C-D) MDS clustering of Geuvadis samples based on proportion of sites with significant AE that differs between sample pairs. Samples are colored by sequencing laboratory and labeled by population. If significant sites are assigned based on a simple binomial test (FDR 5%) the samples cluster first by sequencing laboratory due to lab-specific differences in coverage (c). This effect is mostly removed in (d) by requiring significant sites to have FDR 5% and effect size >0.15.

A challenge in interpreting AE data – whether analyzing binomial p-values or allelic imbalance by other means – are the highly variable total read counts (Fig. 2a), which leads to substantial differences in sampling variance and statistical power between AE sites. This is driven by differences in library depth between samples, as well as biologically variable expression levels between genes and samples. This can affect patterns seen in the data, for example by causing samples to cluster by experimental batch (Fig. 6c) or by tissue (data not shown). If the goal of the analysis is to capture allelic expression, patterns introduced by expression levels are often not desirable. An experimental approach to avoid low read counts in AE data are assays that yield high read counts, such as mmPCR-seq, instead of or alongside with RNA-seq data^9, 12, 13, 40^.

The most conservative method to account for variation in total read counts is to sample total read counts to an even threshold and use these data in downstream analysis. This is a useful approach at least as a sanity check, but vast amounts of valuable data are discarded (Fig. S7c–e). Another straightforward approach is to assign significant sites based on FDR-corrected p-value from the raw counts together with an effect size filter, analogously to differential expression studies – this accounts for the strongest dependency of total read counts (Fig. 6d, S7b). Finally, more sophisticated statistical models can account for various technical sources of variance and integrate information across tissues or individuals to capture phenomena such as *cis*-regulatory variation, nonsense-mediated decay, and imprinting^13, 19^. The most appropriate method depends on the biological question as well as the data type, quality and size, and full benchmarking these approaches is beyond the scope of this paper.

## DISCUSSION

In this paper, we have introduced novel, efficient software tool for retrieving high-quality AE data from RNA-sequencing data sets. We have described how the quality of the input data affects AE analysis, and outlined the quality control approaches that are needed to obtain accurate estimates of allelic expression from RNA-seq data (Fig. S8, Table S1). Altogether, we show that carefully collected and filtered AE estimates from modern RNA-seq data is remarkably robust to technical variation in RNA-sequencing data, highlighting its utility for detecting diverse biological phenomena of genetic and epigenetic variation. Increasingly standardized production of AE data advances wider data sharing and integration across studies, although the geånotype data included in AE estimates by default poses limitations on data access. The increasing size of AE data from large-scale RNA-seq studies hold great promise for capturing regulatory variation even in small numbers of samples, allowing integrated analysis of the personalized genome and its function.

## Acknowledgements

This work was supported by NIH grants 3R01MH101814-02S1, HHSN26820100029C, and 5U01HG006569. We would like to thank the Geuvadis consortium, the GTEx Consortium, the members of the Lappalainen lab, the former GSA group at the Broad, and the bioinformatics team of the New York Genome Center.

## Supplementary Figure Legends

**Figure S1.**
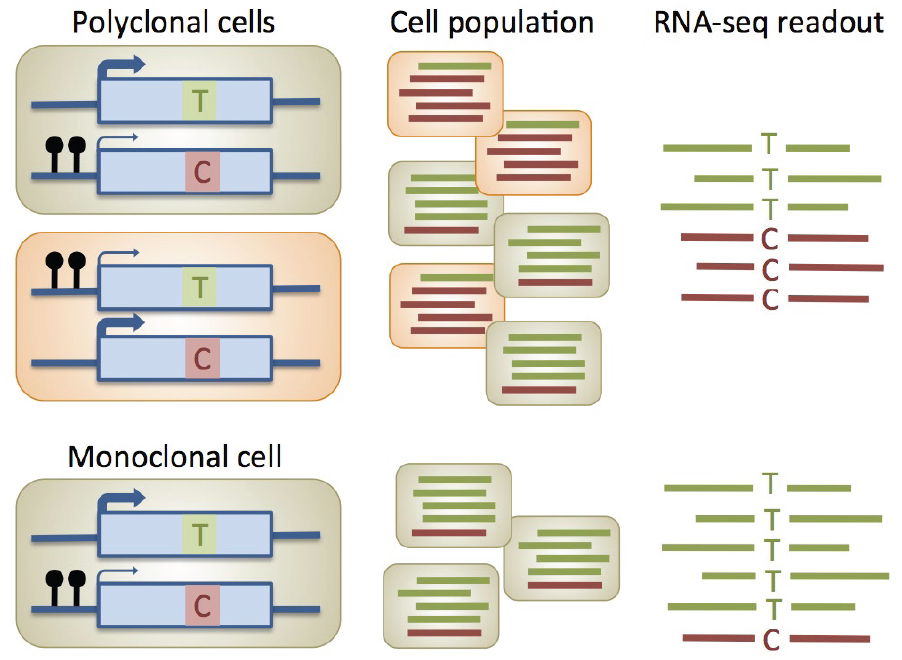
Schematic illustration of allelic expression signal from a population of monoclonal versus polyclonal cells. In the latter, standard RNA-sequencing will show allelic imbalance only when the two alleles are systematically differentially expressed e.g. due to a regulatory variant or imprinting.

**Figure S2.**
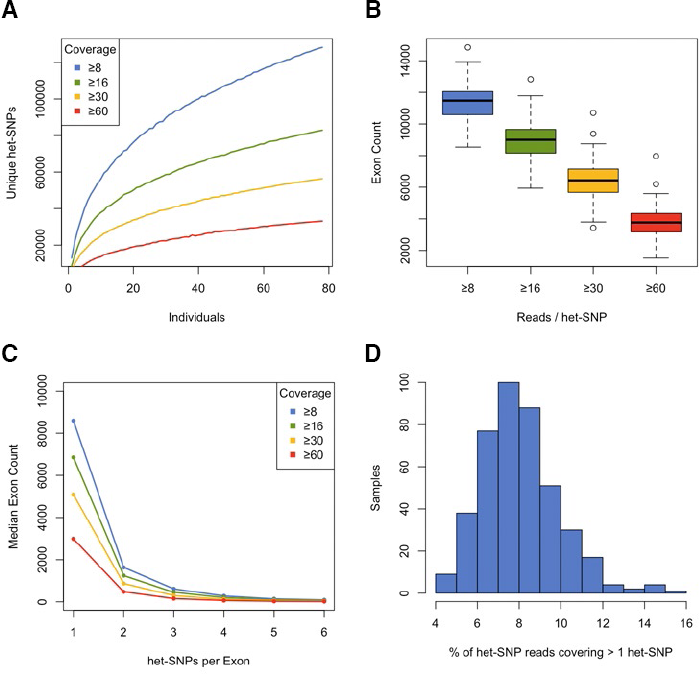
Genomic coverage of allelic expression data in Geuvadis CEU samples (extended). A) Total number of unique het-SNPs covered by increasing read depth as a function of the number of individuals. B) Boxplot of the total number of exons per individual containing at least one het-SNP for each depth level. C) Median number of exons as a function of the number of het-SNPs per feature at increasing read depths. D) Distribution of percentage of reads mapping to het-SNPs that cover > 1 het-SNP for all Geuvadis samples (median = 8.8%).

**Figure S3.**
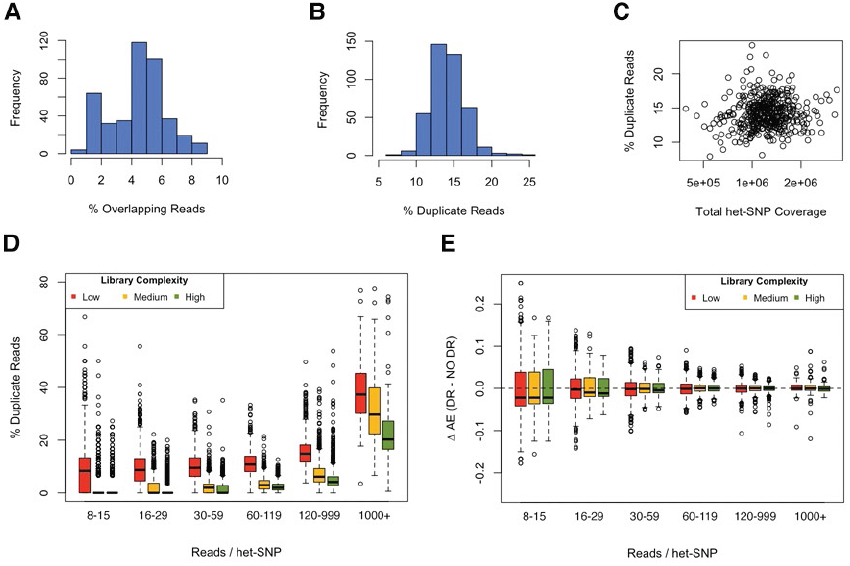
Effect of overlapping and duplicate reads on AE analysis of Geuvadis samples. A) Histogram of percent overlapping mates of paired-end reads at het-SNPs used for AE analysis. B) Histogram of percent duplicate reads at het-SNPs used for AE analysis. C) Total coverage vs percent duplicate reads at AE sites. D) Percent duplicate reads in coverage level bins for Geuvadis samples with the minimum (77.5%, red), median (83.9%, yellow) and maximum (89.6%, green) read complexity at het-SNPs. Complexity is defined as total number of reads mapping to het-SNPs after removing duplicates / number of reads before removing duplicates. E) Effect of duplicate removal on allelic expression effect size (AE = | 0.5 – ref reads / total reads |, ΔAE = AE(Dup Removed) – AE(No Dup Removed)) on het-SNPs binned by coverage level, sites where ΔAE = 0 are not shown.

**Figure S4.**
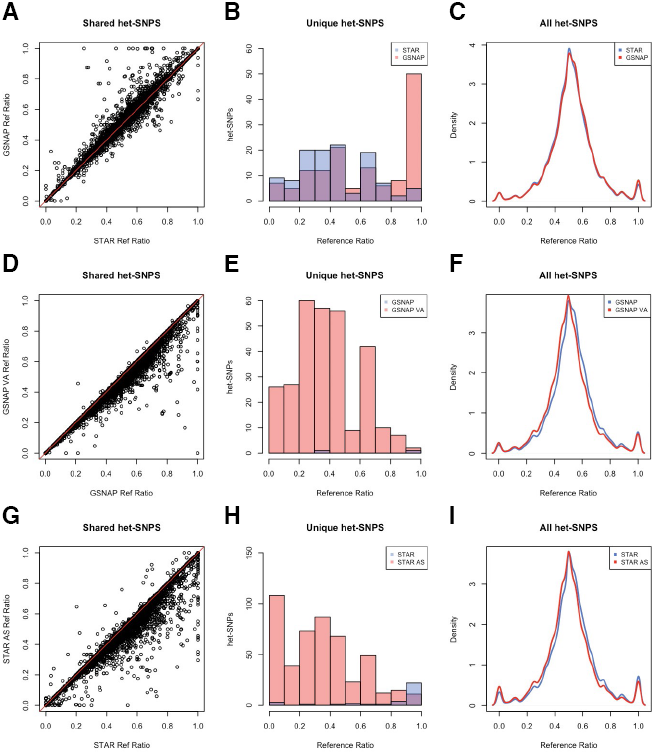
Comparison of AE data generated with different alignment strategies. A-C) Observed reference ratio using STAR 2-pass alignment (STAR) or GSNAP at het-SNPs for which both aligners produce data (a), het-SNPs that only have data using one or the other aligner (b), and the overall distribution of reference ratios (c). D-F) Comparison of GSNAP to GSNAP with variant aware alignment (GSNAP VA), plots as in A-C. G-I) Comparison of STAR mapping to either hg19 or a personalized genome generated with AlleleSeq (STAR AS) as in A-C, without filtering for sites in regions of low mappability or sites that show mapping bias in simulations (see Fig. 3 for analogous filtered data). Sites with low mappability and simulated mapping bias have been excluded from a-f.

**Figure S5.**
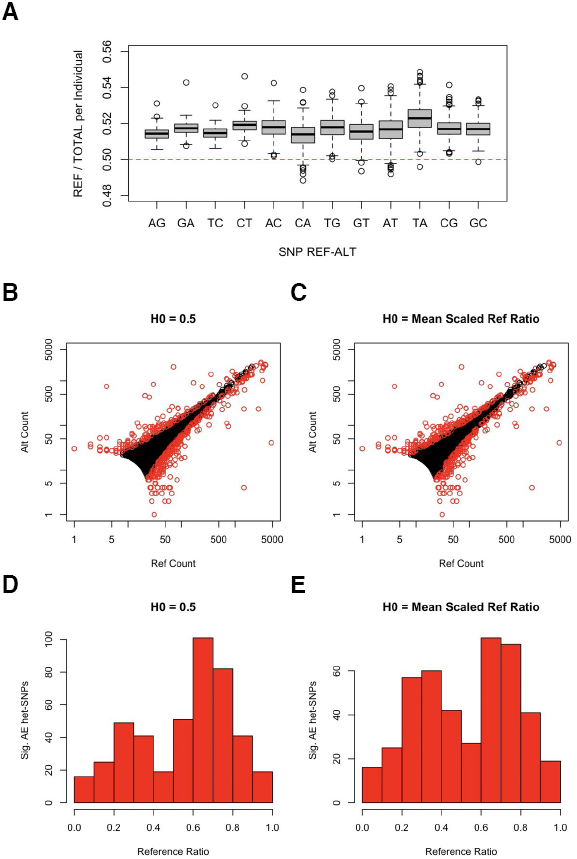
Low-level reference bias at het-SNPs remains after filtering biased sites. A) Boxplot of reference ratio (reference / total) for each reference-alternative base combination for each Geuvadis sample, mapped with STAR 2-pass and filtered for sites with low mappability or mapping bias in simulations as well as sites with potential genotyping error as described before. Ratio is calculated by summing up all REF and ALT read counts for that combination in a sample at sites that have ≥ 8 reads, and for sites with coverage > 75^th^ percentile total counts were scaled down to the 75^th^ percentile to avoid sites with very high coverage having a disproportionate effect on the overall ratio. B-C) Binomial test of AE on an example Geuvadis sample using an expected reference ratio of 0.5 (b) or against the calculated mean scaled reference ratio (c, as described above), with sites of significant AE shown in red (5% FDR). D) Histogram of reference ratios at significant sites from (b), E) Histogram of reference ratios at significant sites from (c).

**Figure S6.**
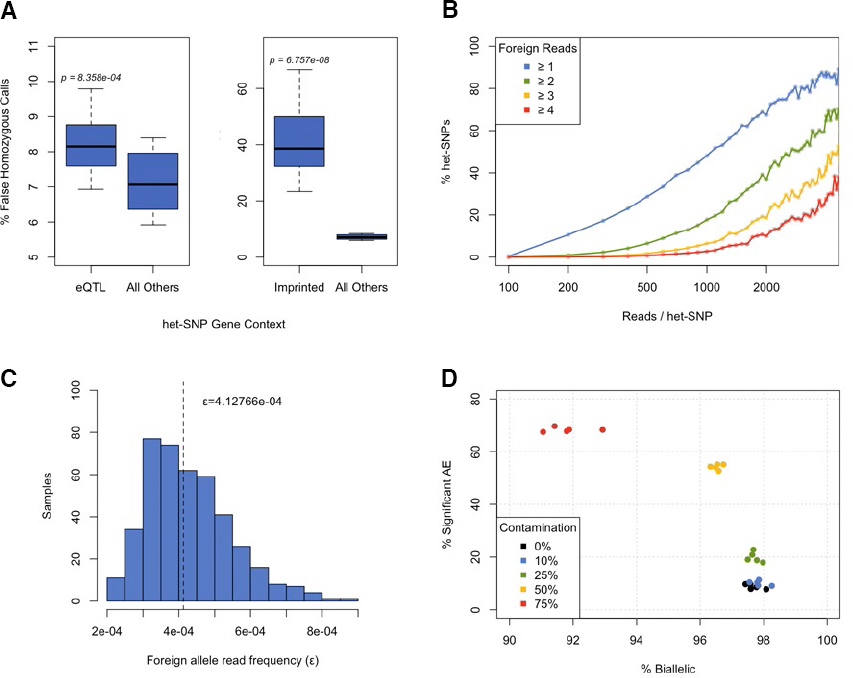
Quality control of genotype data for allelic expression analysis (extended). A) Boxplot of per individual percentage of false homozygous RNA-seq genotype calls at het-SNPs in genes with *cis*-eQTLs in LCLs (FDR <= 0.05, Geuvadis), imprinted genes (based on^13^ excluding genes detected exclusively in Geuvadis data), and all other genes. False homozygosity defined as sites where variant calling using LCL RNA-seq data indicates the individual is homozygous for a non-reference allele, while DNA genotyping (1000 genomes) indicates they are heterozygous. Genotype calls made using GATK and best practices for RNA-seq genotype calling. B) Percentage of het-SNPs where reads from foreign alleles (≥1 blue, ≥2 green, ≥3 yellow, ≥4 red) are observed as a function of coverage level using all Geuvadis RNA-seq data. Binned by hundreds of reads / het-SNP. C) Frequency of the proportion of reads from foreign alleles (non ref or alt) observed (ε) in all Geuvadis samples (median = 4.128 × 10^−4^). D) Scatterplot of percent significant AE sites (binomial test, p < 0.05) and percent biallelic het-SNPs (≥1 read for each allele), for 5 Geuvadis libraries that have been contaminated with another sample *in silico* (0-75% contamination).

**Figure S7.**
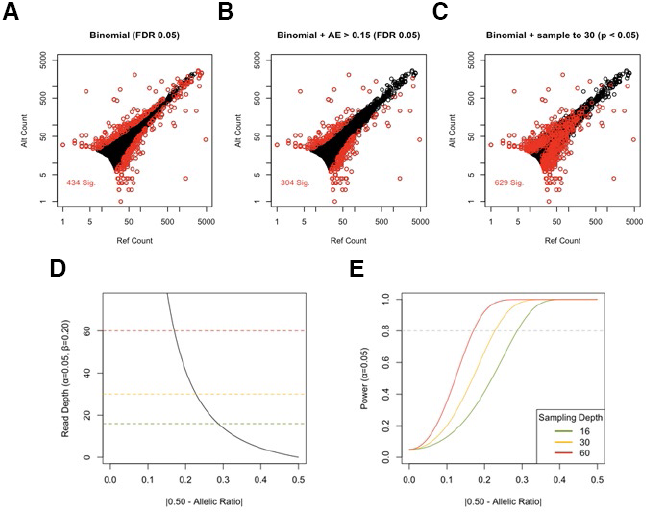
Binomial testing for significant allelic expression (extended). A-C) Testing for significant AE using a binomial test and multiple testing (MT) correction (a, 5% FDR), MT correction and an effect size cutoff (b, 5% FDR, AE), and sampling total het-SNP coverage without replacement to 30 reads with a nominal p-cutoff (c, p < 0.05). D) AE effect size (|0.5 - Ref ratio|) vs statistical power (1 – β) at various sampling depths. E) Minimum effect size that can be detected at a given read depth using a binomial test setting α = 0.05 and β = 0.20.

**Figure S8.**
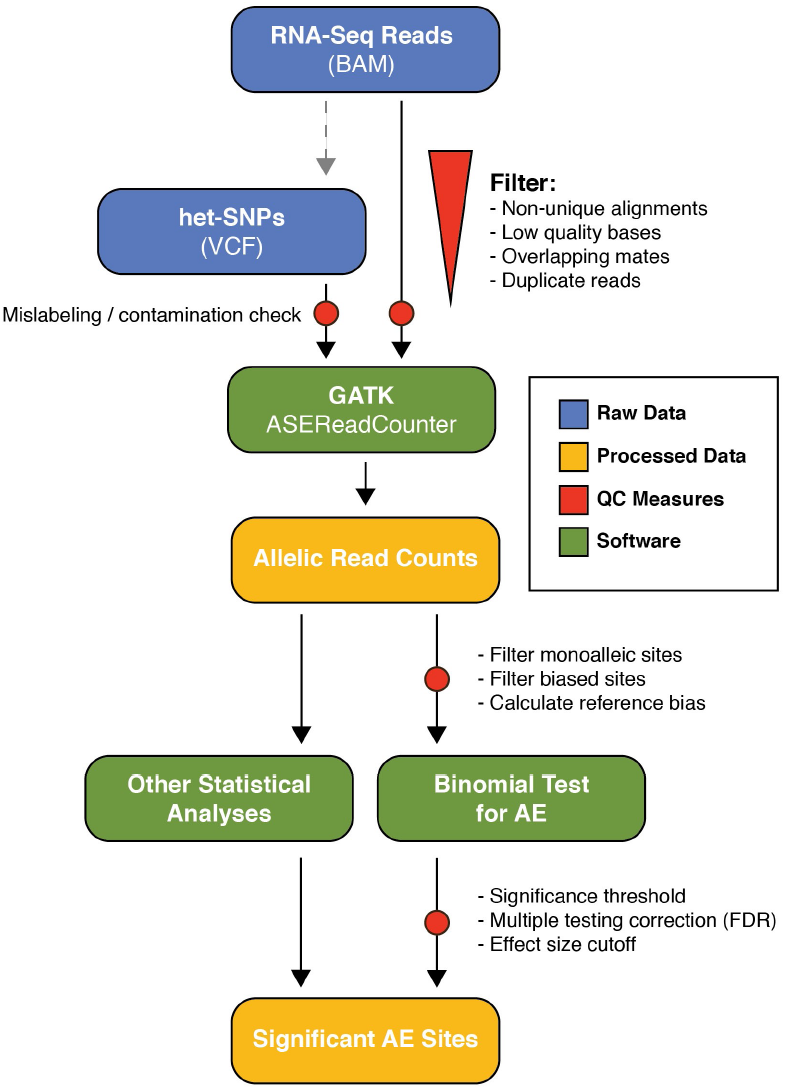
Complete workflow for AE analysis illustrating appropriate quality control measures and filters.

## Supplementary Table Legends

Table S1. Summary of QC problems for AE data, proposed solutions and potential drawbacks.

## ONLINE METHODS

### Filtering homozygous sites

In order to identify potentially homozygous sites miscalled as a heterozygous SNP we model the number of reads that can be observed due to technical error of the experimental and upstream computational pipeline. Let us assume there are a total of *n* reads originating from a site homozygous for an allele R. Assuming a noise rate *ε*, by which a read can erroneously support another allele A, the distribution of total number of reads aligned to allele A, *n*_*A*_, is given by binomial distribution. Hence the probability of a observing *n*_*A*_ or more reads assigned to allele A in a site homozygous for R is given by:

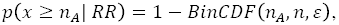

where *BinCDF(n*_*A*_*, n, ε)* is the binomial cumulative distribution function. Conversely, the probability of a observing *n*_*R*_ (n= *n*_*R*_ + *n*_*A*_) or more reads assigned to allele R in a site homozygous for A is given by:

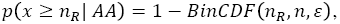

under the assumption that the noise rate is equal for all alleles. Therefore, the probability of observing extreme allelic imbalance due to the null hypothesis, homozygosity for one of the alleles, can be calculated by summing up the two above probabilities corresponding to the two tails of the distribution. In order to derive an empirical estimate of the noise rate *ε* we used ratio between the total sum of reads assigned to other alleles, those different than the designated reference or alternative allele at each site, to the total number of reads in a library divided by two. For this purpose we exclude the sites with more than 5% of the reads aligned to other alleles from the analysis.

### Mapping Strategies for Allelic Expression Analysis

For all analyses, unless otherwise noted, reads were mapped using STAR v2.4.0f1 and the 2-pass mapping strategy as recommended by the Broad (https://www.broadinstitute.org/gatk/guide/article?id=3891). Briefly, splice junctions are detected during a first pass mapping, and these are used to inform a second round of mapping. All reads were mapped to hg19 and Gencode v19 annotations were used.

For mapping to a personalized genome, the X tool, part of AlleleSeq was used to generate both a maternal and paternal genome for NA06986 from the phased 1000 Genomes phase 1 reference using het-SNPs only. Reads were then mapped to both genomes separately using STAR 2-pass strategy (as above). Reads which aligned uniquely to only one genome were kept, and in cases where reads mapped uniquely to both genomes, the alignment with the higher mapping quality was used. The script available for merging alignments to personalized genomes is available online at http://tlab.org.

Mapping using GSNAP was performed with default settings and splice site annotations from hg19 refGene. Variant aware alignment was performed using the “-d” option for NA06986 from the phased 1000 Genomes phase 1 reference using het-SNPs only, as described in the GSNAP documentation.

### MDS Clustering of Samples by Allelic Expression Data

A pairwise distance matrix was produced for all Geuvadis samples using AE data and used for classical multidimensional scaling (cmdscale) in R. The first two dimensions were then plotted against each other for all samples. The distance between two samples was calculated as follows: pairwise distance = total number of sites with significant AE in only one sample / total number of shared sites. A binomial test with a 5% FDR was used for significance with either no effect size cutoff (6c) or a minimum effect size of 0.15 (6d).

### Units of allelic expression

For a single variant:

Reference Ratio = Reference Reads / Total Reads
Allelic Expression (effect size) = | 0.5 – Reference Ratio |

